# Pharmaceutical assessment of low global warming potential alternatives to HFA-134a in a budesonide, glycopyrrolate, and formoterol fumarate pressurized metered dose inhaler

**DOI:** 10.64898/2026.05.12.724523

**Authors:** Kellisa Lachacz, Richard Kaye, Lauren Mello, Ashley Stoker, Johannes Törnell

## Abstract

Manufacturers are adopting propellants for use in pressurized metered-dose inhalers (pMDIs) that have lower global warming potentials (GWPs) than the propellants traditionally used in pMDIs. Hydrofluoroalkane (HFA)-134a has been used as the propellant in the pMDI used to deliver the fixed-dose triple combination of budesonide, glycopyrrolate and formoterol fumarate (BGF); following successful clinical evaluation, the BGF pMDI is now being transitioned to the next generation propellant hydrofluoroolefin (HFO)-1234ze(E), which has near-zero GWP. We describe formulation development efforts that led to selection of HFO-1234ze(E) over another propellant, HFA-152a, for reformulation. Propellant-specific studies evaluated active pharmaceutical ingredient (API) stability and aerodynamic particle size distribution (aPSD). Those analyses have been complemented by *in silico* regional lung deposition modeling conducted after the clinical evaluation of the reformulated BGF pMDI. HFO-1234ze(E) supported favorable stability and aPSD characteristics for BGF pMDI reformulation, compared with HFA-152a, and modeling predicted regional deposition consistent with therapeutic intent. Given that each pMDI is a unique combination of APIs, device, propellant, and excipients, propellant substitution requires product-specific evidence and regulatory approval, and typically takes several years. Targeted analyses, such as those described here, helped to identify the most suitable candidate propellant for successful substitution in the BGF pMDI.

**Highlights:**

- Formulation development efforts that led to evaluation of a budesonide-glycopyrrolate-formoterol fumarate pressurized metered-dose inhaler (BGF pMDI) reformulated with the next generation propellant HFO-1234ze(E) in a clinical trial program are described; the suitability of another propellant, HFA-152a, was also assessed
- Over 6 months under accelerated storage conditions (40°C/75% relative humidity [RH]), the HFA-152a formulation approached and, in one replicate, fell below the 90% of formulation label claim threshold of evaluation, whereas the original HFA-134a product and the HFO-1234ze(E) formulation remained above that threshold
- Over 6 months under accelerated storage conditions (40°C/75% RH) and 18 months under long-term stability storage conditions (25°C/60% RH), the fine particle mass and fine particle fraction for all active pharmaceutical ingredients (APIs) showed that the HFO-1234ze(E) formulation tracked more closely than the HFA-152a formulation to the original HFA-134a product
- Later *in silico* modeling, conducted after clinical testing, predicted a trend for greater deposition of APIs in early airway generations with HFA-152a, whereas HFO-1234ze(E) was predicted to more closely match HFA-134a, indicating a greater likelihood of achieving equivalence to the original HFA-134a product with HFO-1234ze(E) than with HFA-152a
- Based on these analyses and other formulation development efforts, HFO-1234ze(E) was identified as the most suitable propellant for reformulation of the BGF pMDI; for HFA-152a, analyses raised concerns about storage stability, and differences in aerosol characteristics that can impact API deposition in the lungs and, in turn, efficacy

## Introduction

Pressurized metered-dose inhalers (pMDIs) are used by millions of people with chronic obstructive pulmonary disease (COPD) and asthma, including many elderly people, young children and people with low inspiratory flow, for whom dry-powder inhalers are not clinically suitable.^1,2^ Propellant selection for pMDIs is important in ensuring and optimizing storage stability and therapeutic performance.^3^ Although medical aerosols account for less than 0.04% of total greenhouse gas emissions,^4^ the growing threat of climate change means that there is a need to reduce that burden and the global warming potential (GWP) of propellants used in pMDIs has therefore become an important consideration.

*Breztri Aerosphere/Trixeo Aerosphere* is a single-inhaler, fixed-dose triple combination of budesonide, glycopyrrolate, and formoterol fumarate (BGF) delivered by a pMDI. It is approved for the treatment of patients with COPD in more than 80 countries, including the USA,^5^ and in the European Union (for patients with moderate to severe COPD not adequately treated with dual therapy).^6^ It is currently the only triple combination medicine approved for the treatment of patients with COPD that uses this specific combination of active pharmaceutical ingredients (APIs), each of which has properties that affect efficacy, safety and formulation. Hereafter, it is referred to as ‘the BGF pMDI’. Historically, the propellant used in the BGF pMDI has been hydrofluoroalkane (HFA)-134a (1,1,1,2-tetrafluoroethane). However, the GWP of HFA-134a, on a 100-year horizon, is 1300 times greater than that of CO_2_^7^ and it is subject to phase-out under the European F-Gas Regulation, and phase-down under the Kigali Amendment to the Montreal Protocol or the American Innovation and Manufacturing Act (AIM).^8-10^ Options to replace HFA-134a as the propellant used in the BGF pMDI, with a propellant with lower GWP, were therefore explored.

Given the complexity of pMDI formulations, with each being a unique combination of APIs, device, propellant and other excipients, propellant substitution requires comprehensive testing and success is uncertain. As well as ensuring pharmaceutical quality is maintained (e.g. physiochemical storage stability and dosing consistency), a novel propellant requires toxicological assessment and demonstration of *in vitro* aerodynamic particle size distribution (aPSD) and clinical bioequivalence to allow transfer of existing regulatory approvals without repeating phase 3 clinical trials of efficacy and safety.

Limitations were identified with some potential candidate propellants to replace HFA-134a, including the toxicity of hydrofluoroolefin (HFO)-1234yf (2,3,3,3-tetrafluoropropene), the flammability of dimethyl ether and saturated light hydrocarbons, and the unacceptable physical and material properties of compressed gases (e.g. air, CO_2_, nitrogen).^11^ Only two candidate propellants were considered suitable for the initial pharmaceutical assessment to replace HFA-134a in the BGF pMDI : HFO-1234ze(E) (1,3,3,3-tetrafluoropropene), and HFA-152a (1,1-difluoroethane), despite the flammability of the latter.^11^ The GWPs of HFO-1234ze(E) and of HFA-152a, on 100-year horizons, are 99.9% lower and approximately 90% lower than the GWP of HFA-134a, respectively.^7^ Like HFA-134a, HFA-152a is subject to phase-out under the European F-Gas Regulation, or phase-down under the Kigali Amendment to the Montreal Protocol or AIM, whereas HFO-1234ze(E) is not.^8-10^

*In vitro* equivalence of the BGF pMDI formulated with HFO-1234ze(E) to the original HFA-134a product has been demonstrated across delivered dose uniformity, fine particle mass, spray pattern and plume geometry.^12^ Subsequently, the BGF pMDI formulated with HFO-1234ze(E) was found to meet bioequivalence criteria with the original HFA-134a product across all APIs in healthy adults, in a phase 1 randomized controlled trial.^13^ Furthermore, the HFO-1234ze(E) formulation was found to have similar safety profiles to the original HFA-134a product over 12 and 52 weeks, with no new safety signals, in patients with COPD in a phase 3 randomized controlled trial.^14^ The BGF pMDI is now being transitioned from the original HFA-134a product to the HFO-1234ze(E) formulation in routine clinical practice where it has received regulatory approval, including in the European Union,^6^ to reduce the greenhouse gas emissions associated with its use while ensuring its continued availability.

The objective of this report is to summarize additional key data from pharmaceutical assessments of stability under storage and *in vitro* aerosol properties, which informed the decision to evaluate the BGF pMDI formulated with HFO-1234ze(E) in a clinical trial program. More recently generated *in silico* regional lung deposition prediction data are also presented.

## Experimental

### Assessments of active pharmaceutical ingredient stability and aerodynamic particle size distribution

#### Materials and sample preparation

Co-suspensions of the APIs in individual BGF pMDI canisters were prepared by suspending spray-dried lipid porous particles with micronized crystalline drug particles in propellants (HFA-134a, HFO-1234ze(E) or HFA-152a). The preparation of the porous particles and pMDI filling process have been described previously.^15,16^ The same processes were used to prepare sample canisters containing each propellant. The three formulations were designed to deliver the same mass of porous particles and drug particles in each actuation across the three propellants. The suspensions were transferred into 14 mL fluorinated ethylene polymer-coated aluminum canisters (Presspart, Blackburn, UK) through 50 μL valves (Bespak, King’s Lynn, UK), which are standard pMDI components.

#### Active pharmaceutical ingredient stability and degradation products assays

The concentrations of each API and their corresponding degradation products in the sample canisters were determined by reversed-phase high-performance liquid chromatography (HPLC) with ultra-violet detection. Sample canisters were frozen, their valves were cut off, and their contents were transferred to 50 mL centrifuge tubes, allowing the propellant to evaporate completely. The formulations’ solids were extracted and analyzed via HPLC. The net fill weights of the canisters were determined. The concentration of each API was calculated using the quantified concentration of the API divided by the net fill weight. The concentrations were reported as a percent of the label claim for each formulation. The concentration of the degradation products was calculated by dividing the peak area of the degradation product in the chromatogram by the peak area of the corresponding API. The concentrations of the degradation products were reported as a percent of the corresponding API concentration.

Stability was assessed over 6 months under accelerated storage conditions of 40°C/75% relative humidity (RH) and over 18 months under the climatic zone II long-term stability conditions (25°C/60% RH), with inhalers in a valve-downwards orientation and protected in an overwrap with a desiccant sachet. Stability of each formulation was assessed according to prespecified thresholds.

#### Aerodynamic particle size distribution

aPSD of each API was determined with HFA-134a, HFO-1234ze(E) or HFA-152a using a Next Generation Impactor (NGI) operated at 30 L/min, as described in United States Pharmacopeia (USP), chapter 601.^17^ Sample canisters were placed into actuators with two waste priming actuations and two additional waste seating actuations. Five actuations were collected in the NGI with a USP induction port attached. The valve, actuator, induction port, and NGI collection cups were extracted with volumetrically dispensed solvent. The sample solutions were assayed using a drug-specific chromatographic method. The fine particle mass, < 5 μm, was defined as the sum of API masses collected on the impactor stages with aerodynamic cut-off diameters below 5.0 μm at the test flow rate. Testing was conducted over 6 months under accelerated storage conditions (40°C/75% RH) and over 18 months under long-term stability conditions (25°C/60% RH).

#### In silico *prediction modeling of regional lung deposition*

The regional deposition model is based on that described by the National Council on Radiation Protection and Measurements in 1997,^18^ applied to the Yeh and Schum lung structure model.^19^ It represents filters-in-series where for each airway generation the deposition fraction of each particle size bin is calculated from the probabilities of deposition by impaction, sedimentation and Brownian diffusion. Deposition is predicted using an exaggerated breath-hold time of 30 seconds,^20^ practically ignoring deposition in the exhalation phase; this is consistent with experimentally determined exhaled fractions for the BGF pMDI, which are less than 0.5% irrespective of breath-hold time.^21^ Data were binned by groupings of airway generations corresponding to bronchi (generations 0–8), bronchioles (9–16) and the alveolar region (> 16).

## Results

### Active pharmaceutical ingredient stability and degradation products assays

Over 6 months under accelerated storage conditions (40°C/75% RH), all three propellant formulations showed a reduction in formoterol fumarate as a percent of the formulations’ label claim (**Figure 1**). The HFA-152a formulation approached and, in one replicate at 6 months, fell below the 90% threshold of evaluation, whereas the HFA-134a and HFO-1234ze(E) formulations remained within the 90%–100% thresholds of evaluation over 6 months. For all three propellants tested, in line with expectations, the proportion of formoterol fumarate degradation products increased over 6 months, with the rate of increase being greatest with HFA-152a (**Figure 2**). All formulations remained within the thresholds of evaluation for the glycopyrrolate and budesonide assays, with no unexpected increase in degradation products, over 6 months under the accelerated storage conditions (**Supplemental Figures 1–4**). Over 18 months under long-term stability conditions (25°C/60% RH) all formulations remained within the 90%–110% thresholds for all APIs, with no unexpected increases in degradation products (**Supplemental Figures 5–10**).

**Figure 1.**
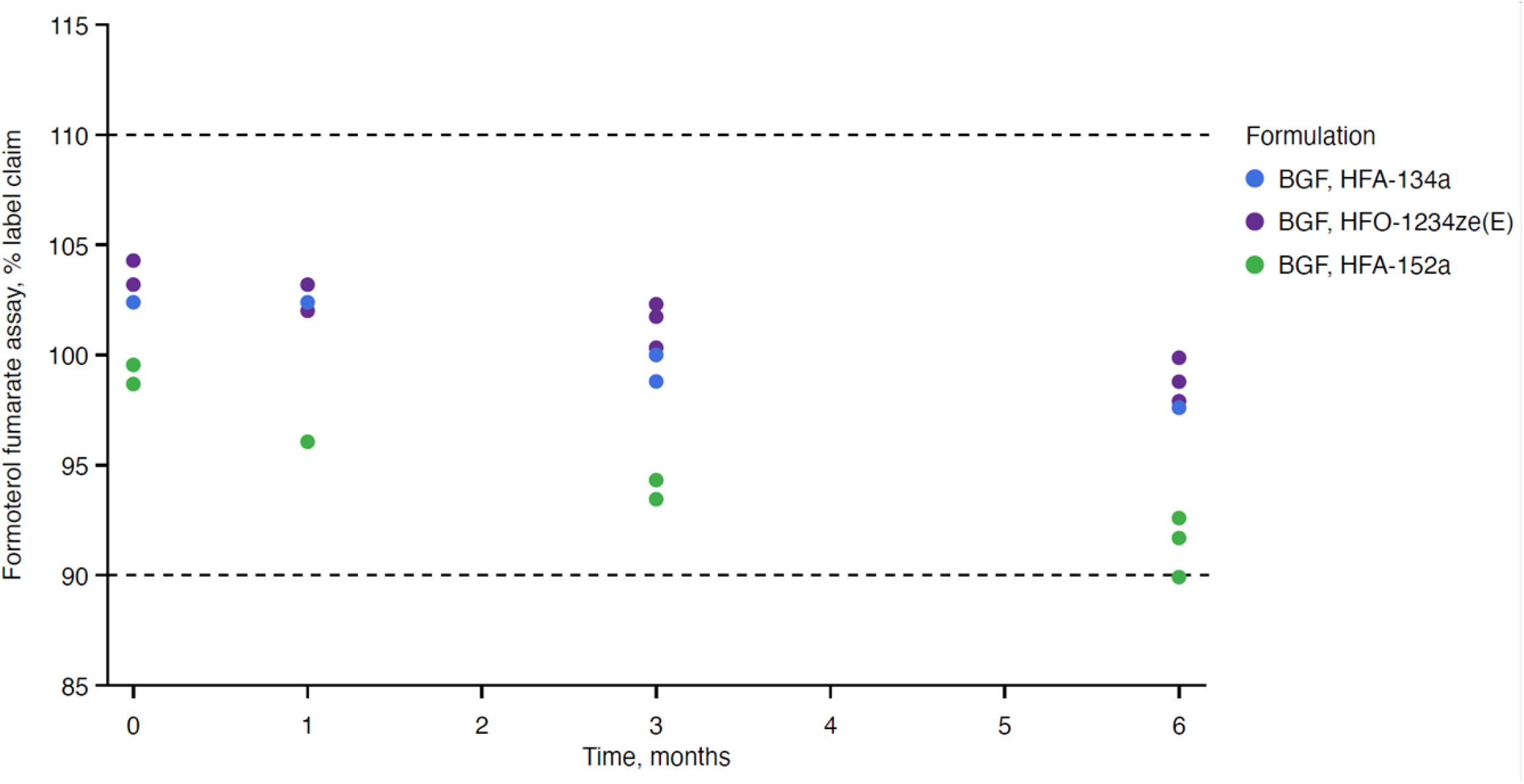
Stability: assay of formoterol fumarate, expressed as a percent of the label claim, in each propellant under accelerated stability conditions (40°C/75% RH) over 6 months. Dashed lines represent the threshold of evaluation at 90% and 110% of the formulations’ label claim. BGF, budesonide, glycopyrrolate and formoterol fumarate; HFA, hydrofluoroalkane; HFO, hydrofluoroolefin; RH, relative humidity

**Figure 2.**
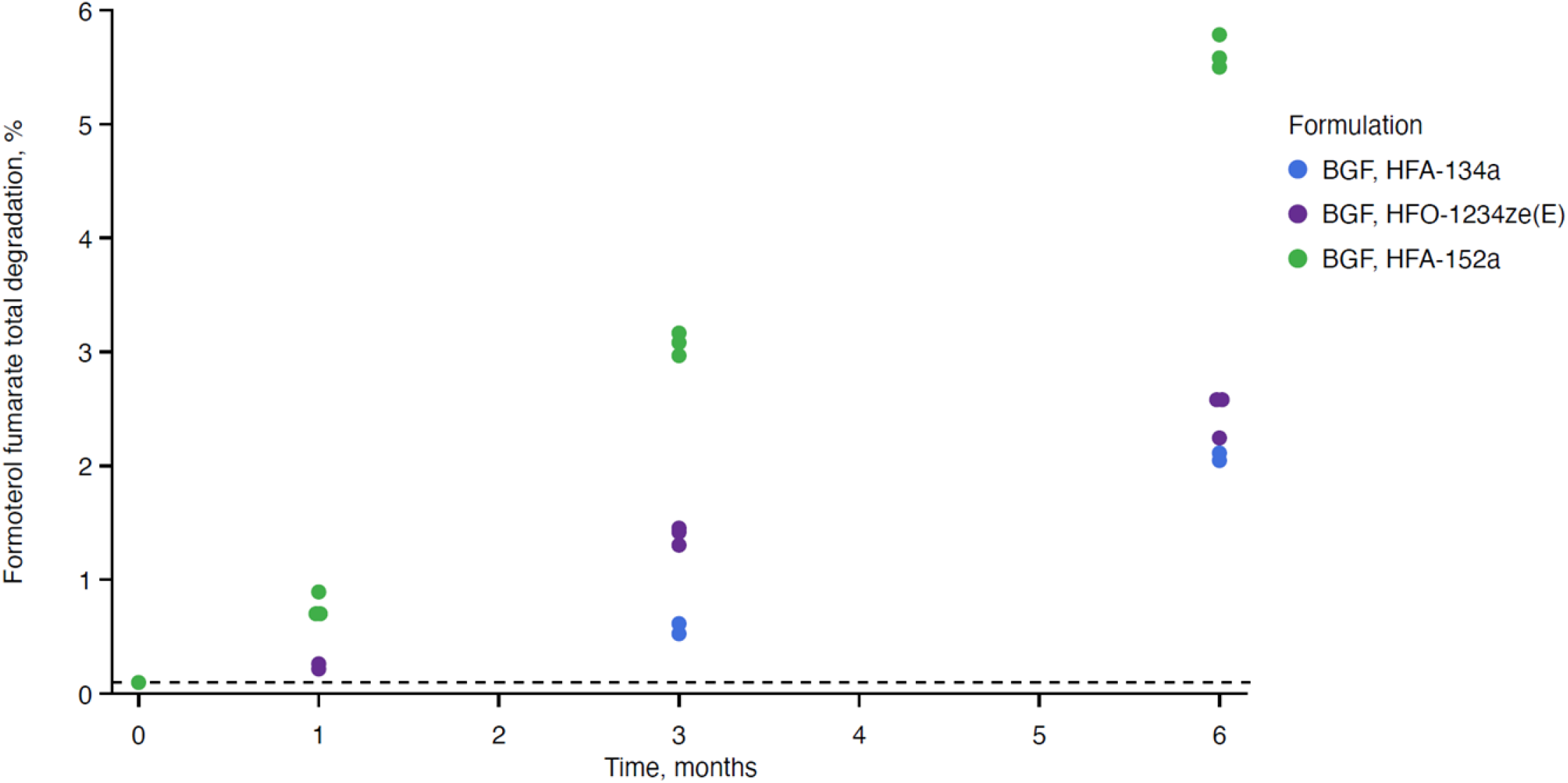
Stability: total degradation products for formoterol fumarate in each propellant under accelerated stability conditions (40°C/75% RH) over 6 months. Dotted line represents the limit of quantification, 0.1%. BGF, budesonide, glycopyrrolate and formoterol fumarate; HFA, hydrofluoroalkane; HFO, hydrofluoroolefin; RH, relative humidity

### Aerodynamic particle size distribution

Fine particle mass < 5μm, expressed as micrograms/actuation, of formoterol fumarate was lower in the HFA-152a formulation than in the original HFA-134a product and the HFO-1234ze(E) formulation, under the accelerated storage conditions (40°C/75% RH) over 6 months (**Figure 3**). Similar results were seen for glycopyrrolate and budesonide (**Supplemental Figures 11 and 12**). Over 18 months under long-term stability conditions (25°C/60% RH), fine particle mass of all APIs was also lower, across most measurements, with HFA-152a than with HFA-134a or with HFO-1234ze(E) (**Supplemental Figures 13– 15**). For fine particle fraction, expressed as a percent of the label claim, the HFO-1234ze(E) formulation more closely approximated to the original HFA-134a product, across both storage conditions tested and all APIs, than the HFA-152a formulation (**Supplemental Figures 16 and 17**).

**Figure 3.**
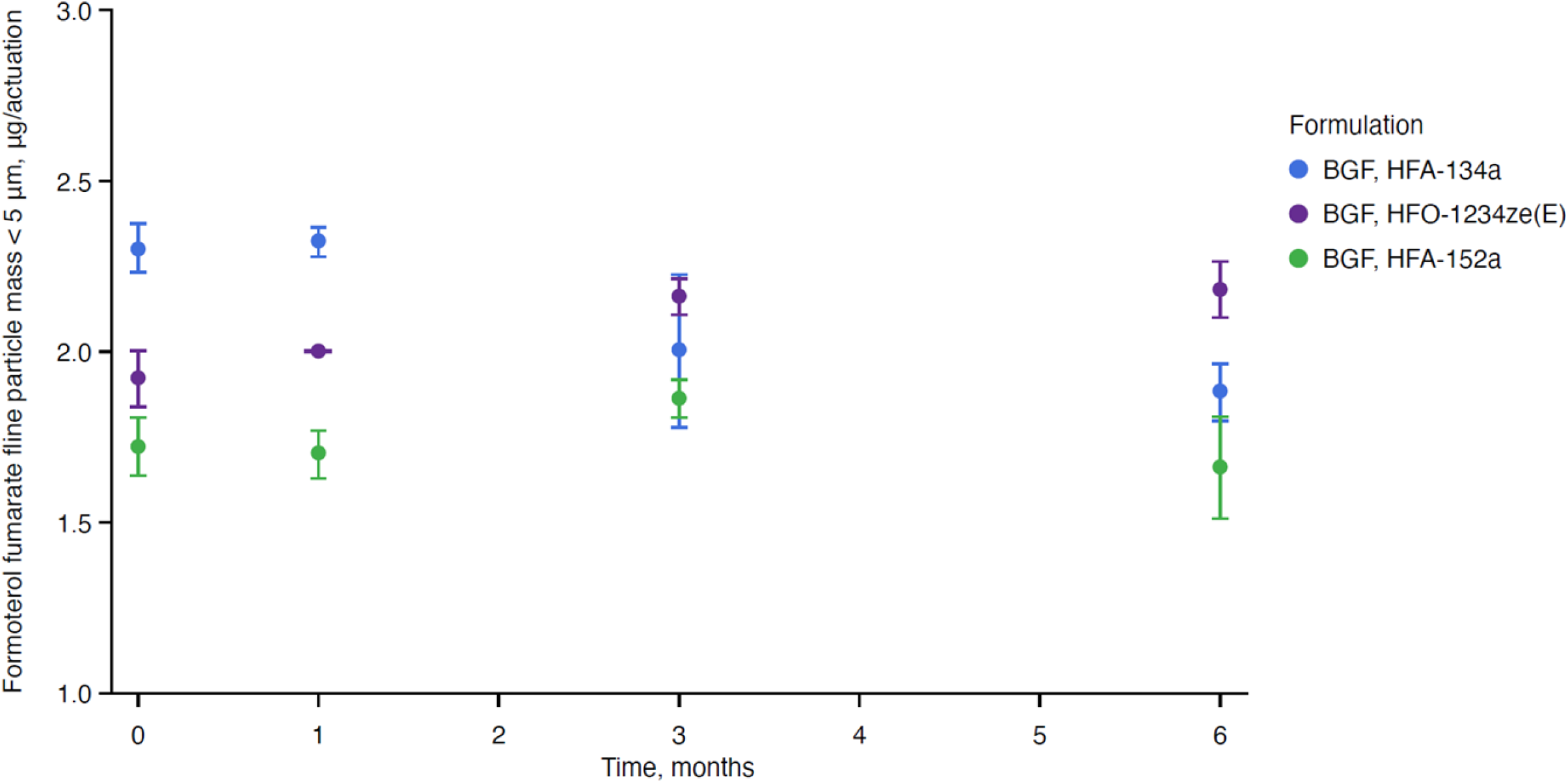
Aerosol: fine particle mass < 5μm, expressed as micrograms/actuation, for formoterol fumarate in each propellant under accelerated stability conditions (40°C/75% RH) over 6 months. Results are presented as mean ± standard deviation BGF, budesonide, glycopyrrolate and formoterol fumarate; HFA, hydrofluoroalkane; HFO, hydrofluoroolefin; RH, relative humidity

### In silico *prediction modeling of regional lung deposition*

Compared with HFA-152a, HFO-1234ze(E) is predicted to more closely match the BGF lung deposition of the original HFA-134a product (**Figure 4**). With HFA-152a, there is a predicted trend for greater deposition in the bronchi (airway generations 0–8) and less deposition in the alveolar region (airway generations > 16) than with HFA-134a or with HFO-1234ze(E).

**Figure 4.**
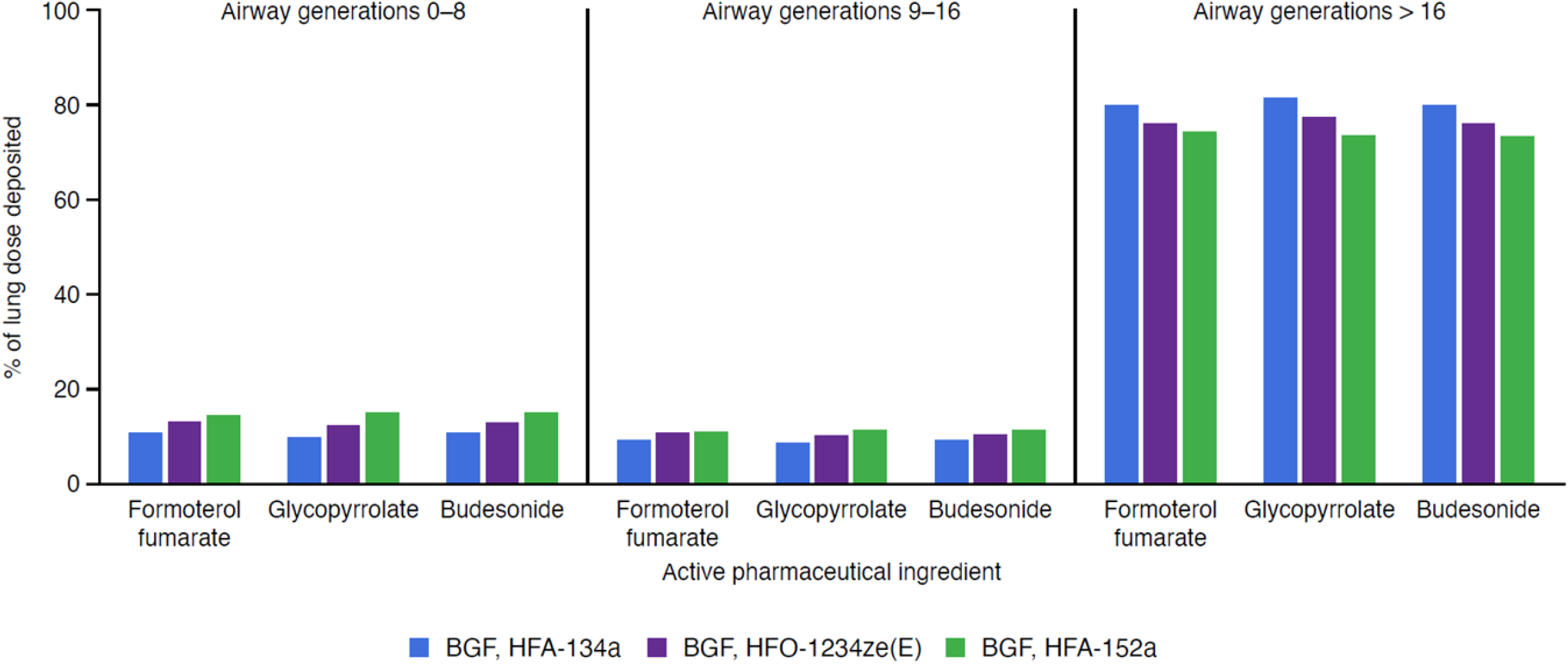
Regional lung deposition: budesonide, glycopyrrolate and formoterol fumarate (in silico modeling) BGF, budesonide, glycopyrrolate and formoterol fumarate; HFA, hydrofluoroalkane; HFO, hydrofluoroolefin

## Discussion

The results of the analyses presented here indicated that HFO-1234ze(E) may be a suitable substitute for HFA-134a in the BGF pMDI, as was later demonstrated in further preclinical and clinical testing.^12-14^ For HFA-152a, the analyses indicated a substantial risk of non-substitutability. Data that most clearly indicated potential substitutability issues have been addressed directly; however, all results from the studies described are available in the Supporting Information.

Concerns identified about the potential substitutability of HFA-152a in the BGF pMDI included chemical instability, and greater differences in *in vitro* aerosol characteristics between the HFA-152a formulation and the original HFA-134a product, compared with the HFO-1234ze(E) formulation. Aerosol characteristics are crucial to ensuring that APIs reach their sites of action in the peripheral airways;^22^ any differences in API deposition resulting from changes in these characteristics could affect efficacy.^23^

Concerns about API stability related particularly to formoterol fumarate, which is a sensitive API with well-characterized degradation pathways (e.g. hydrolysis).^24^ It is plausible that the greater extent and rate of degradation of formoterol fumarate with HFA-152a, compared with HFA-134a and HFO-1234ze(E) under an accelerated storage condition, reflects the higher water solubility of HFA-152a. Compared with formoterol fumarate, budesonide and glycopyrrolate are both more resistant to degradation.

Based on these analyses, the BGF pMDI formulated with HFA-152a would have a reduced shelf life compared to the HFA-134a product, possibly requiring refrigerated storage. The risks of reformulating with HFA-152a, as a viable commercial product, were greater than reformulating than with HFO-1234ze(E) due to the potential need for a refrigerated supply chain, concerns about shelf life and the flammability of HFA-152a^11^ potentially requiring extensive adaptation of manufacturing facilities. This was one important factor in the decision to focus reformulation efforts on HFO-1234ze(E). In other analyses relating to shelf life, although not to substitutability, we found higher leak rates with the standard commercial valve in the BGF pMDI formulated with HFO-1234ze(E) or with HFA-152a than in the original HFA-134a product; valve optimization was subsequently conducted for HFO-1234ze(E) formulation, but was not conducted for HFA-152a.

Differences in fine particle mass and fine particle fraction among the three propellant formulations tested were seen across all APIs. The *in vitro* NGI analyses found that the proportion of fine particles was generally lower with HFA-152a than with HFA-134a, with the HFO-1234ze(E) formulation aerosol characteristics being more similar to those of the original HFA-134a product. This is in line with the physical characteristics of HFA-152a, which is known to deliver larger droplets than HFA-134a.^25^

*In silico* regional lung deposition modeling predicts that the BGF pMDI formulated with HFA-152a would trend toward earlier deposition in the respiratory tract than with HFA-134a or HFO-1234ze(E). If different proportions of the APIs reach different airway generations within the lungs that may lead to the reformulated treatment not achieving clinical bioequivalence with the original product. Although no such testing of clinical bioequivalence has been undertaken for the BGF pMDI formulated with HFA-152a, these analyses further support the decision that was taken to focus on reformulation with HFO-1234ze(E).

Propellant substitution for a pMDI requires years of work and significant investment. Formulation development efforts, of the types described here, are important in maximizing the likelihood of a successful substitution and informed the decision to reformulate the BGF pMDI with HFO-1234ze(E) rather than HFA-152a. We note that a reformulation with HFA-152a of another pMDI platform that is used to deliver a different fixed-dose triple combination of a different combination of APIs (beclomethasone, glycopyrrolate and formoterol fumarate), which is used for maintenance treatment of patients with COPD or asthma,^26^ has recently undergone evaluation of long-term safety and tolerability in a phase 3 randomized controlled trial.^27^ However, as reflected by regulatory requirements for approval, every pMDI / pMDI platform is different, in terms of APIs, excipients, devices and how they work together, and any changes require specific assessment.

Based on our formulation development efforts, including those presented here, the NGP HFO-1234ze(E) was considered the best candidate for reformulation of the BGF pMDI. Reformulation with HFA-152a presented substantial challenges, with key areas of concern including storage stability and differences in aerosol characteristics that can impact API deposition in the lungs and, in turn, efficacy.

## Abbreviations

AIM: American Innovation and Manufacturing
API: active pharmaceutical ingredient
aPSD: aerodynamic particle size distribution
BGF: budesonide, glycopyrrolate and formoterol fumarate
COPD: chronic obstructive pulmonary disease
GWP: global warming potential
HFA: hydrofluoroalkane
HFO: hydrofluoroolefin
HPLC: high-performance liquid chromatography
LGWP: low global warming potential
NGI: Next Generation Impactor
pMDI: pressurized metered-dose inhaler
RH: relative humidity
USP: United States Pharmacopeia

## Supporting Information

The supplementary figures associated with this article can be found in the online version.

## Acknowledgments

The authors are grateful to Amie Baker, Dafni Bika, Tobias Bramer, Katharine Knappenberger, Ekaterina Maslova, Richard Maughan, Joseph Muscat and Roshy Pakdaman for valuable discussions and insightful comments, which helped greatly in development of this manuscript. Medical writing support was provided by Luke Bratton and Colin Glen of Oxford PharmaGenesis, Oxford, UK, and was funded by AstraZeneca.

## Author contributions

Kellisa Lachacz, Lauren Mello and Ashley Stoker made substantial contributions to the conception and design of the analyses of active pharmaceutical ingredient stability and aerodynamic particle size distribution, and to acquisition, analysis and interpretation of the data.

Richard Kaye made substantial contributions to the conception and design of the *in silico* modeling of regional lung deposition, and to interpretation of the data. Johannes Törnell made substantial contributions to the acquisition, analysis and interpretation of the modeling data.

All authors contributed to drafting and/or reviewed drafts critically for important intellectual content and provided final approval of the version to be published.

