## Supporting Information for "Pharmaceutical assessment of low global warming potential alternatives to HFA-134a in a budesonide, glycopyrrolate, and formoterol fumarate pressurized metered dose inhaler"

Supplemental figure 1. Stability: assay of glycopyrrolate, expressed as a percent of the label claim, in each propellant under accelerated stability conditions (40°C/75% RH) over 6 months

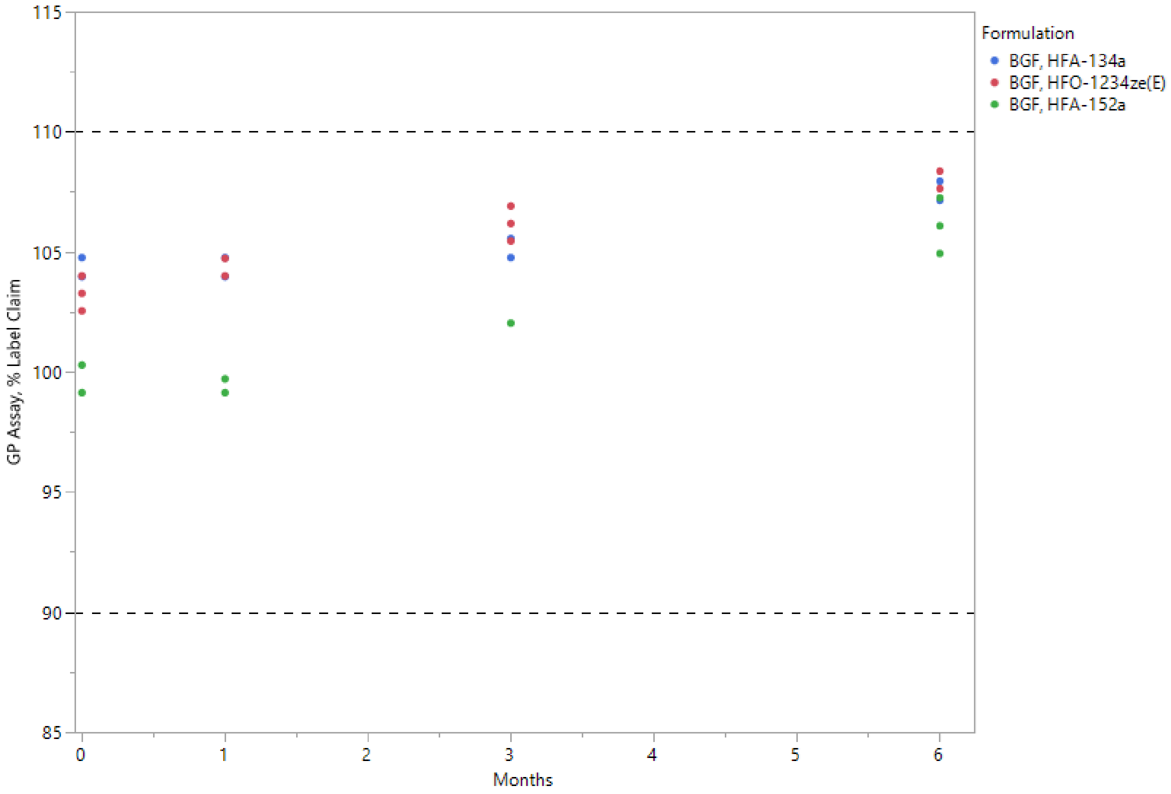


Dashed lines represent the threshold of evaluation at 90% and 110% of the formulations' label claim.

BGF, budesonide, glycopyrrolate and formoterol fumarate; GP, glycopyrrolate; HFA, hydrofluoroalkane; HFO, hydrofluoroolefin; RH, relative humidity

Supplemental figure 2. Stability: assay of budesonide, expressed as a percent of the label claim, in each propellant under accelerated stability conditions (40°C/75% RH) over 6 months

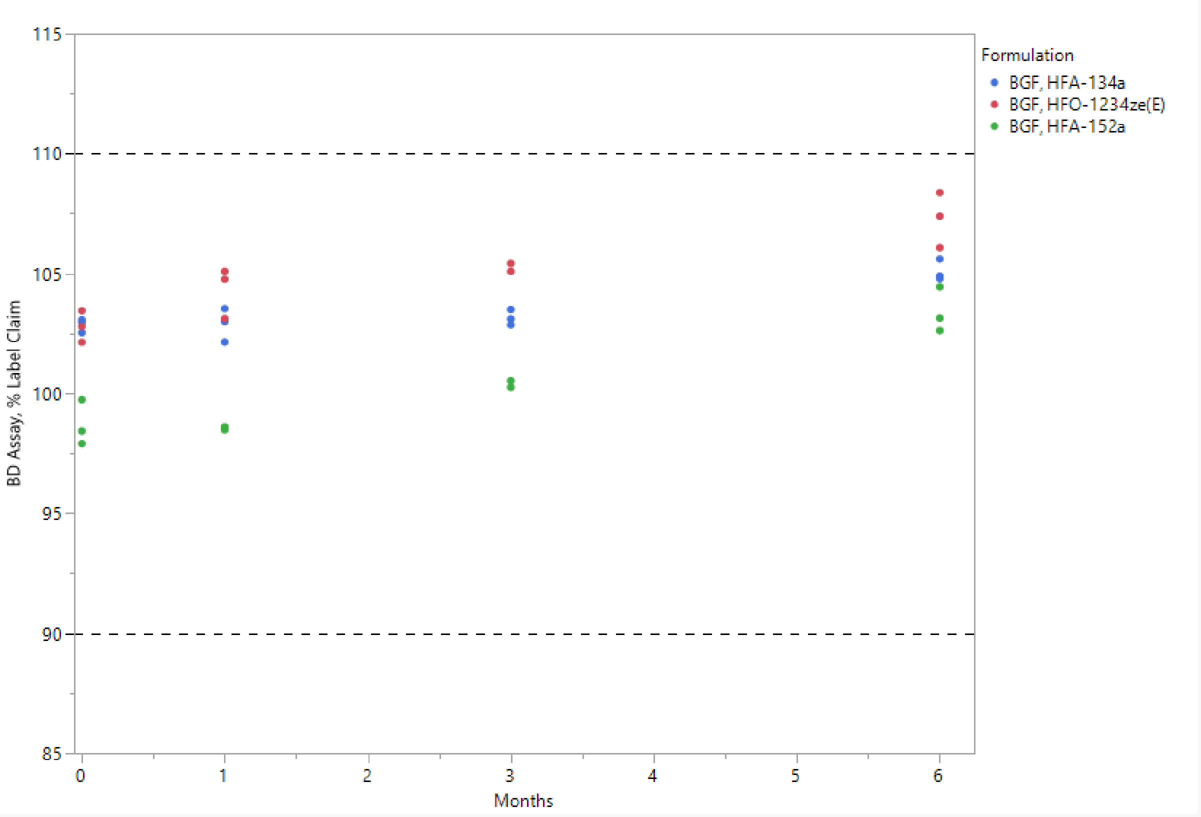


Dashed lines represent the threshold of evaluation at 90% and 110% of the formulations' label claim.

BD, budesonide; BGF, budesonide, glycopyrrolate and formoterol fumarate; HFA, hydrofluoroalkane; HFO, hydrofluoroolefin; RH, relative humidity

Supplemental figure 3. Stability: total degradation products for glycopyrrolate in each propellant under accelerated stability conditions (40°C/75% RH) over 6 months

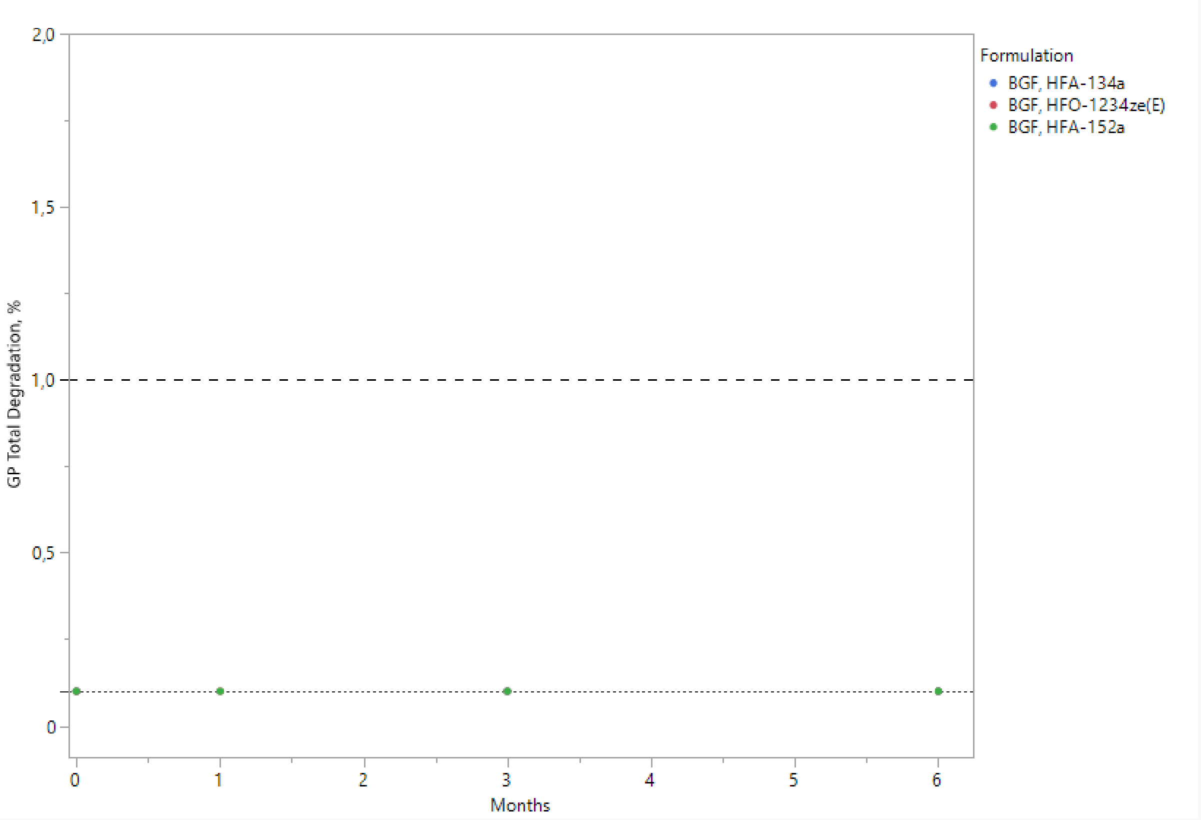


The dotted line represents the limit of quantification, 0.1%.

The dashed line represents the ICH threshold of evaluation,1.0%.

Results for all three propellants were below the limit of quantification.

BGF, budesonide, glycopyrrolate and formoterol fumarate; GP, glycopyrrolate; HFA, hydrofluoroalkane; HFO, hydrofluoroolefin; ICH, International Council for Harmonisation of Technical Requirements for Pharmaceuticals for Human Use; RH, relative humidity

Supplemental figure 4. Stability: total degradation products for budesonide in each propellant under accelerated stability conditions (40°C/75% RH) over 6 months

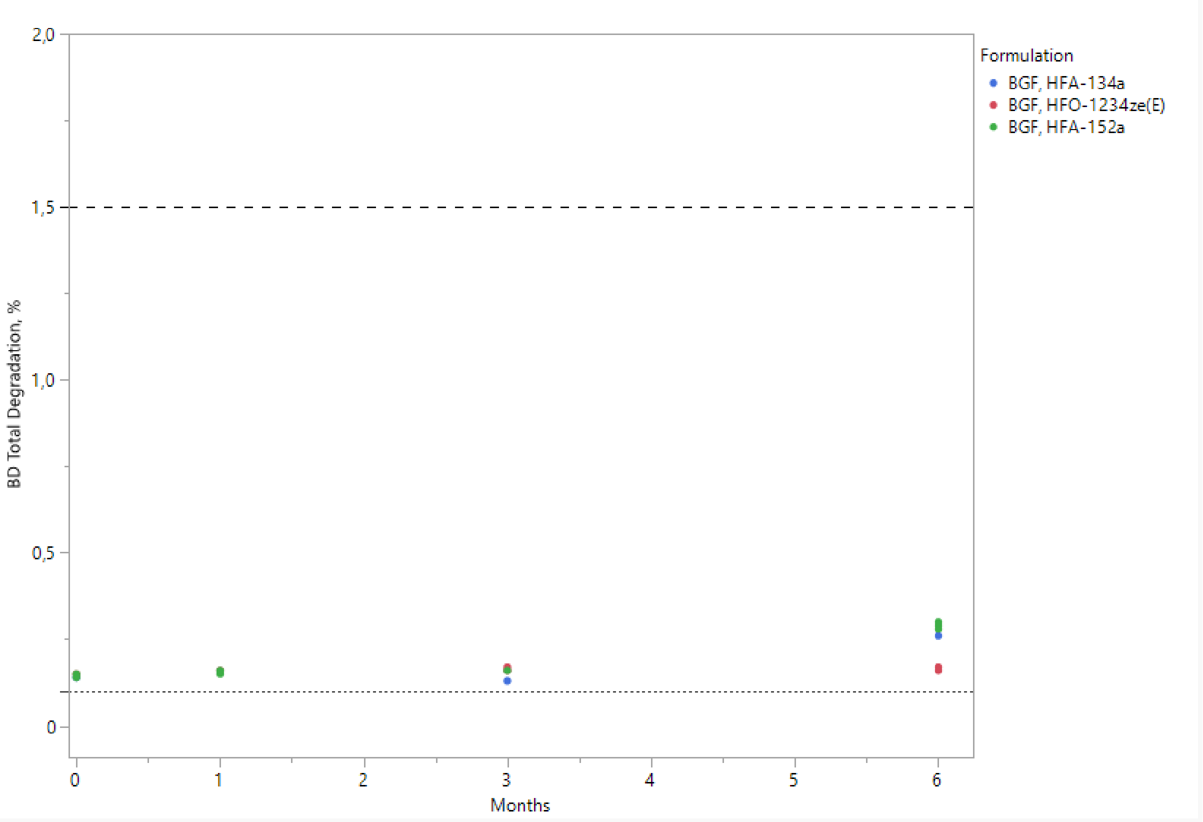


Dotted line represents the limit of quantification, 0.1%.

The dashed line represents the threshold of evaluation,1.5%, per the BP monograph for budesonide pressurized inhalation.

BD, budesonide; BGF, budesonide, glycopyrrolate and formoterol fumarate; BP, British Pharmacopoeia; HFA, hydrofluoroalkane; HFO, hydrofluoroolefin; RH, relative humidity

Supplemental figure 5. Stability: assay of formoterol fumarate, expressed as a percent of the label claim, in each propellant under long-term stability conditions (25°C/60% RH) over 18 months

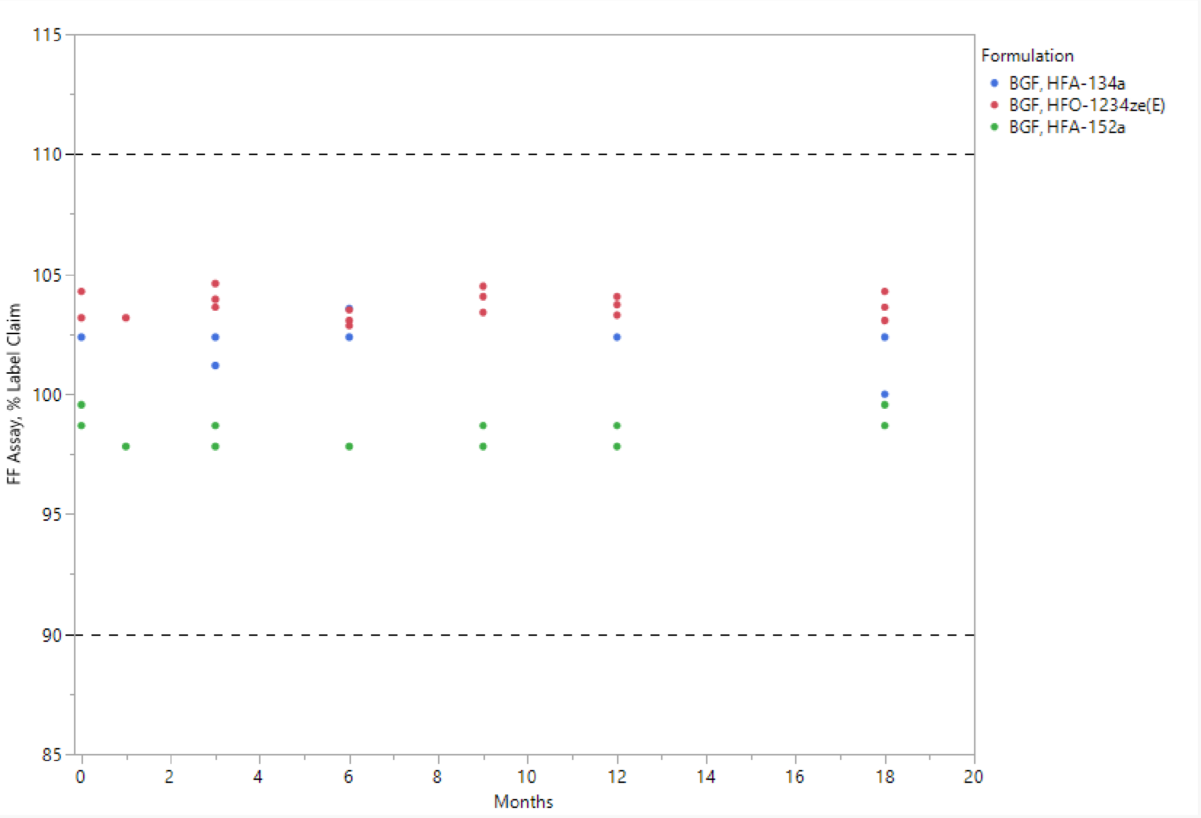


Dashed lines represent the threshold of evaluation at 90% and 110% of the formulations' label claim.

BGF, budesonide, glycopyrrolate and formoterol fumarate; FF, formoterol fumarate; HFA, hydrofluoroalkane; HFO, hydrofluoroolefin; RH, relative humidity

Supplemental figure 6. Stability: assay of glycopyrrolate, expressed as a percent of the label claim, in each propellant under long-term stability conditions (25°C/60% RH) over 18 months

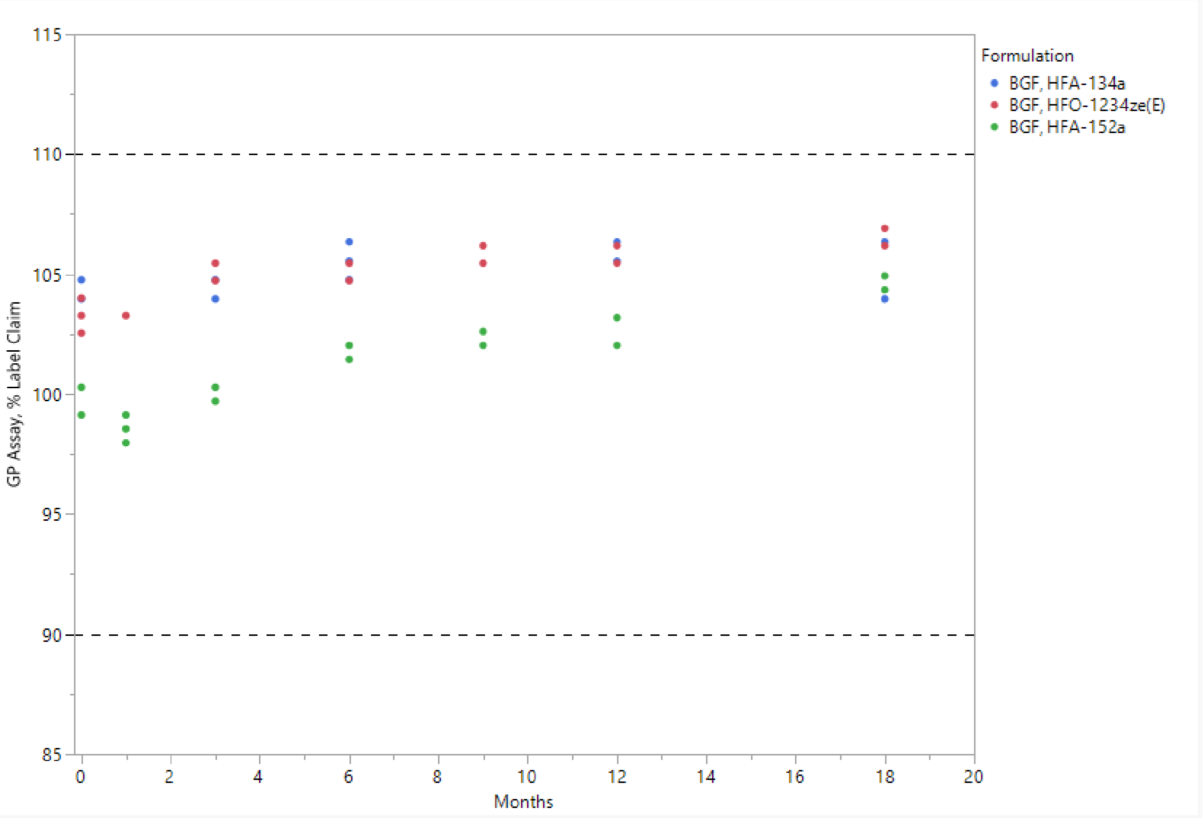


Dashed lines represent the threshold of evaluation at 90% and 110% of the formulations' label claim.

BGF, budesonide, glycopyrrolate and formoterol fumarate; GP, glycopyrrolate; HFA, hydrofluoroalkane; HFO, hydrofluoroolefin; RH, relative humidity

Supplemental figure 7. Stability: assay of budesonide, expressed as a percent of the label claim, in each propellant under long-term stability conditions (25°C/60% RH) over 18 months

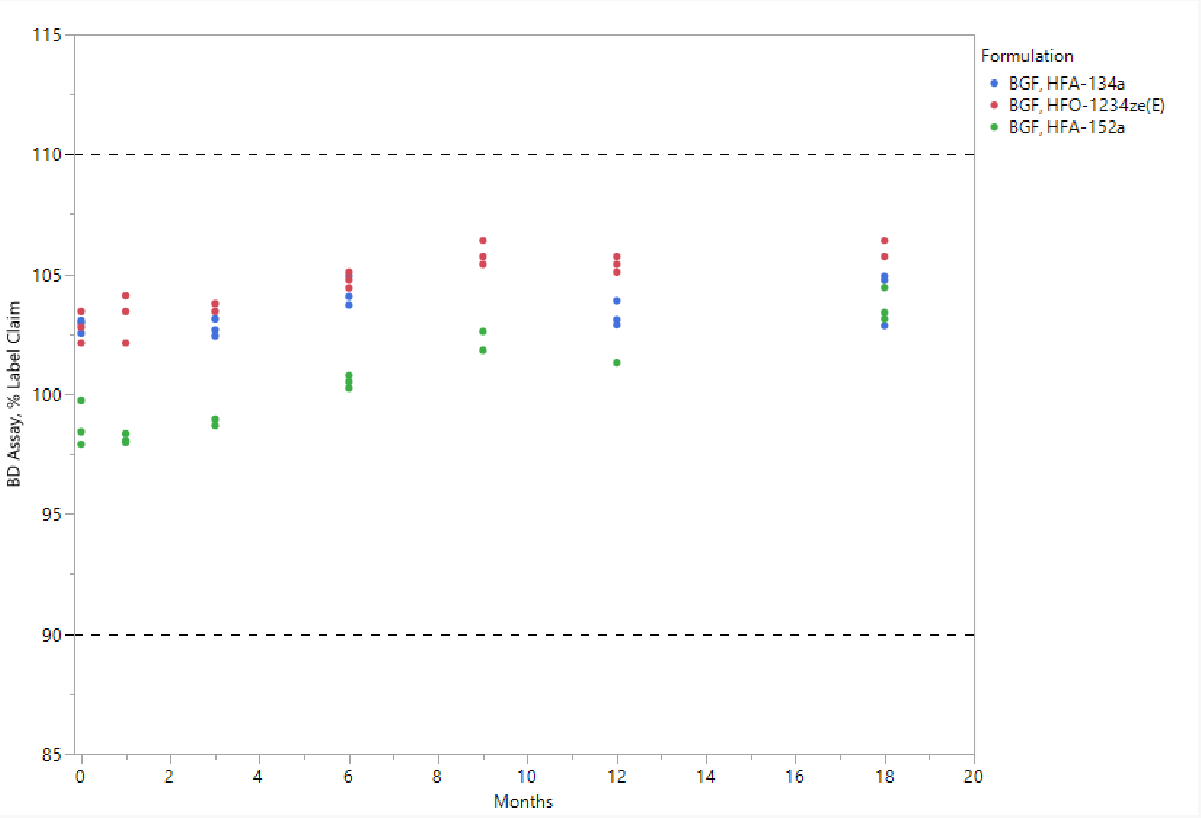


Dashed lines represent the threshold of evaluation at 90% and 110% of the formulations' label claim.

BD, budesonide; BGF, budesonide, glycopyrrolate and formoterol fumarate; HFA, hydrofluoroalkane; HFO, hydrofluoroolefin; RH, relative humidity

Supplemental figure 8. Stability: total degradation products for formoterol fumarate in each propellant under long-term stability conditions (25°C/60% RH) over 18 months

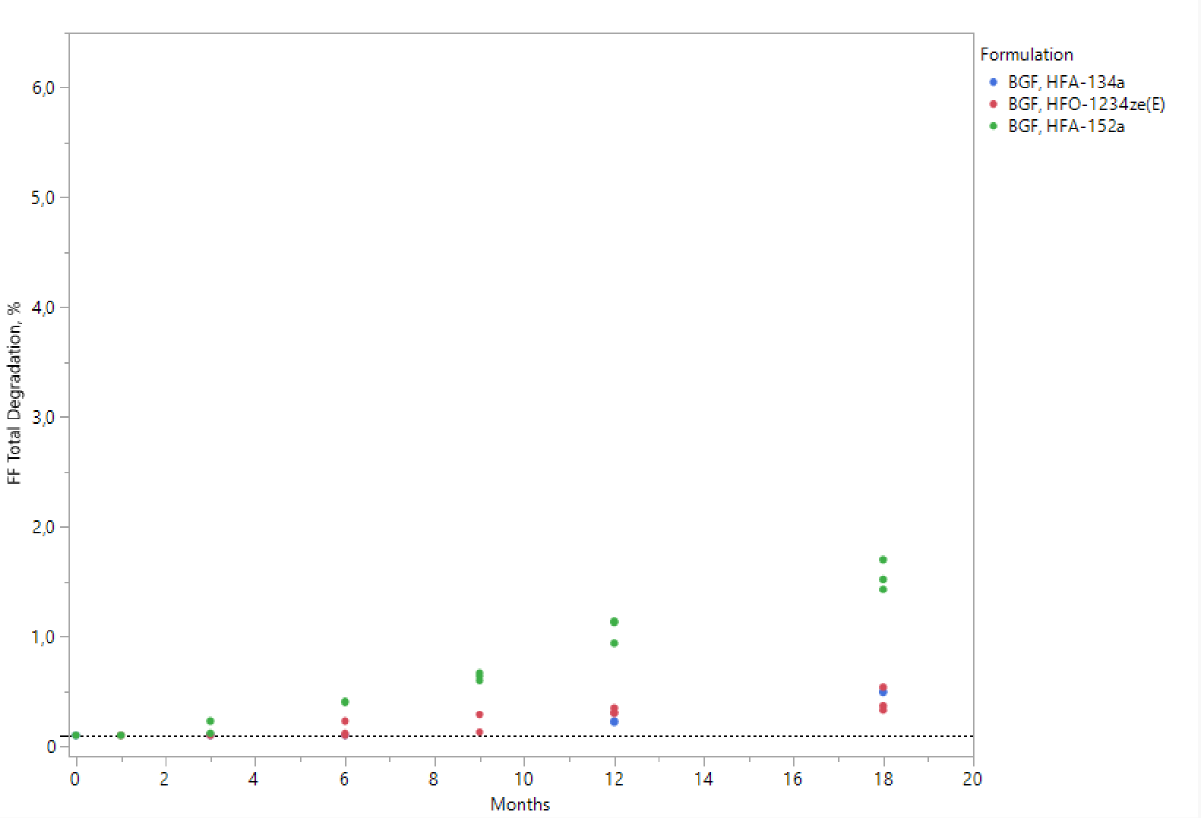


The dotted line represents the limit of quantification, 0.1%.

BGF, budesonide, glycopyrrolate and formoterol fumarate; FF, formoterol fumarate; HFA, hydrofluoroalkane; HFO, hydrofluoroolefin; RH, relative humidity

Supplemental figure 9. Stability: total degradation products for glycopyrrolate in each propellant under long-term stability conditions (25°C/60% RH) over 18 months

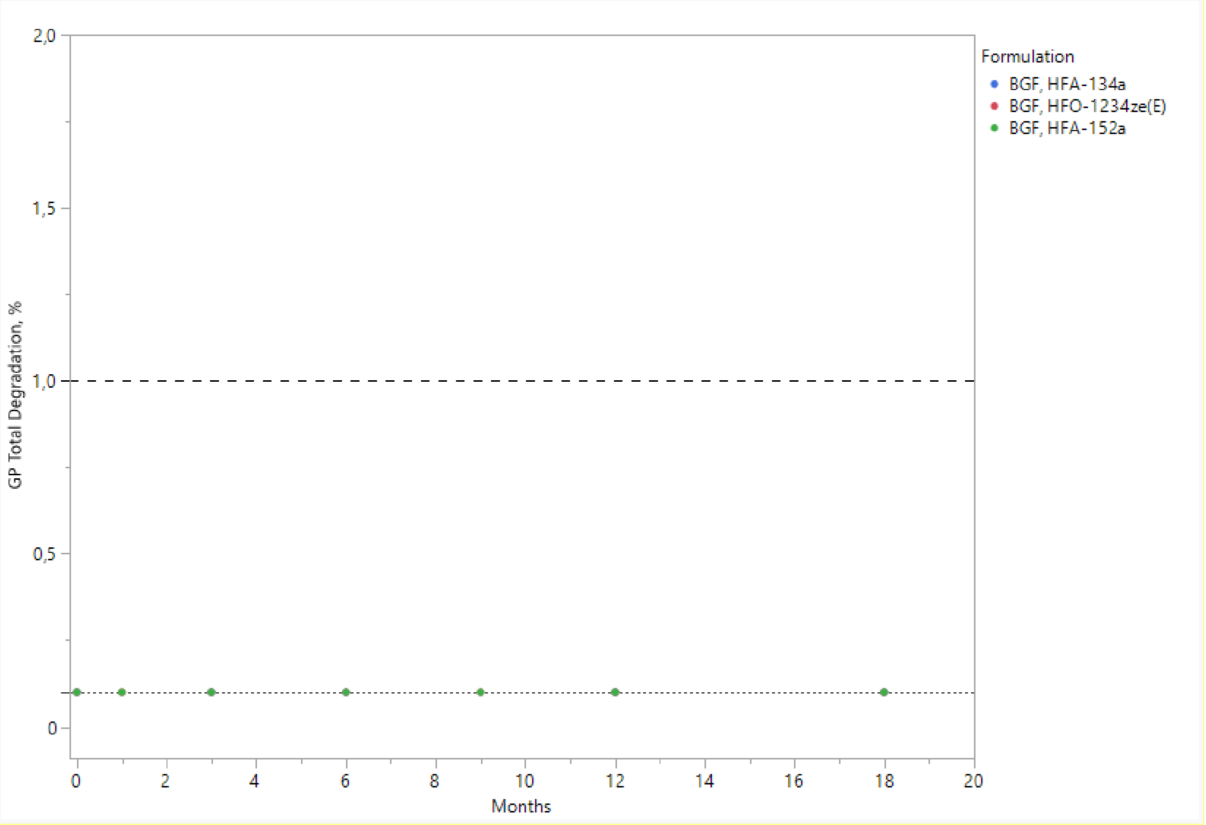


The dotted line represents the limit of quantification, 0.1%.

The dashed line represents the ICH threshold of evaluation,1.0%.

Results for all three propellants were below the limit of quantification.

BGF, budesonide, glycopyrrolate and formoterol fumarate; GP, glycopyrrolate; HFA, hydrofluoroalkane; HFO, hydrofluoroolefin; ICH, International Council for Harmonisation of Technical Requirements for Pharmaceuticals for Human Use; RH, relative humidity

Supplemental figure 10. Stability: total degradation products for budesonide in each propellant under long-term stability conditions (25°C/60% RH) over 18 months

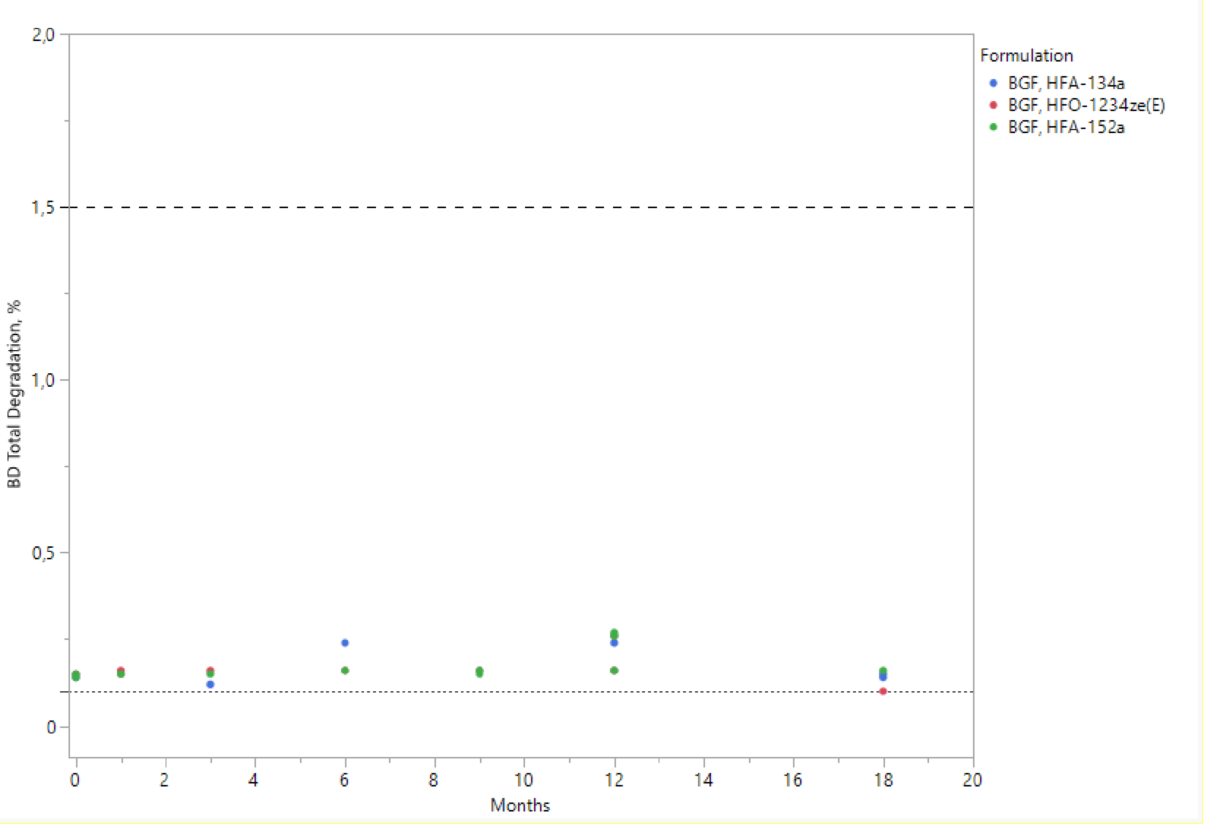


Dotted line represents the limit of quantification, 0.1%.

The dashed line represents the threshold of evaluation,1.5%, per the BP monograph for budesonide pressurized inhalation.

BD, budesonide; BGF, budesonide, glycopyrrolate and formoterol fumarate; BP, British Pharmacopoeia; HFA, hydrofluoroalkane; HFO, hydrofluoroolefin; RH, relative humidity

Supplemental figure 11. Aerosol: fine particle mass < 5μm, expressed as micrograms/actuation, for glycopyrrolate in each propellant under accelerated stability conditions (40°C/75% RH) over 6 months

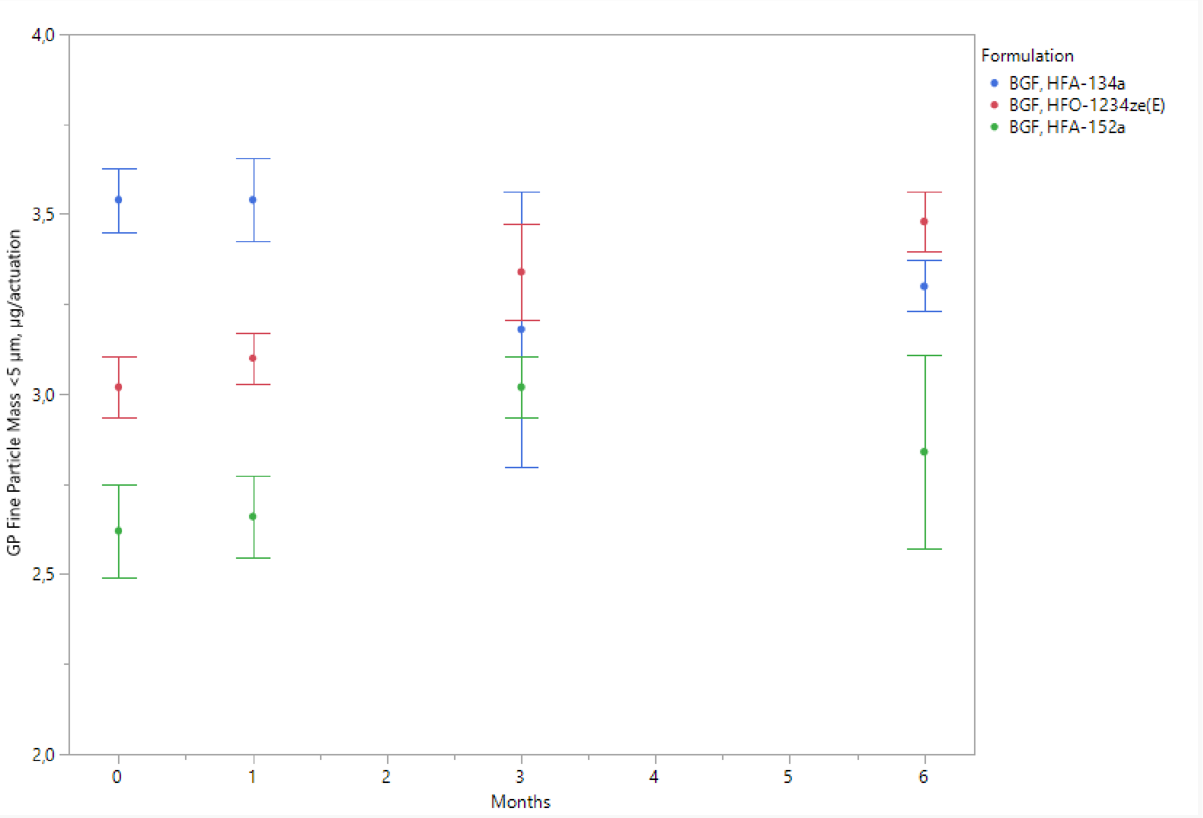


Results are presented as mean ± standard deviation.

BGF, budesonide, glycopyrrolate and formoterol fumarate; GP, glycopyrrolate; HFA, hydrofluoroalkane; HFO, hydrofluoroolefin; RH, relative humidity

Supplemental figure 12. Aerosol: fine particle mass < 5µm, expressed as micrograms/actuation, for budesonide in each propellant under accelerated stability conditions (40°C/75% RH) over 6 months

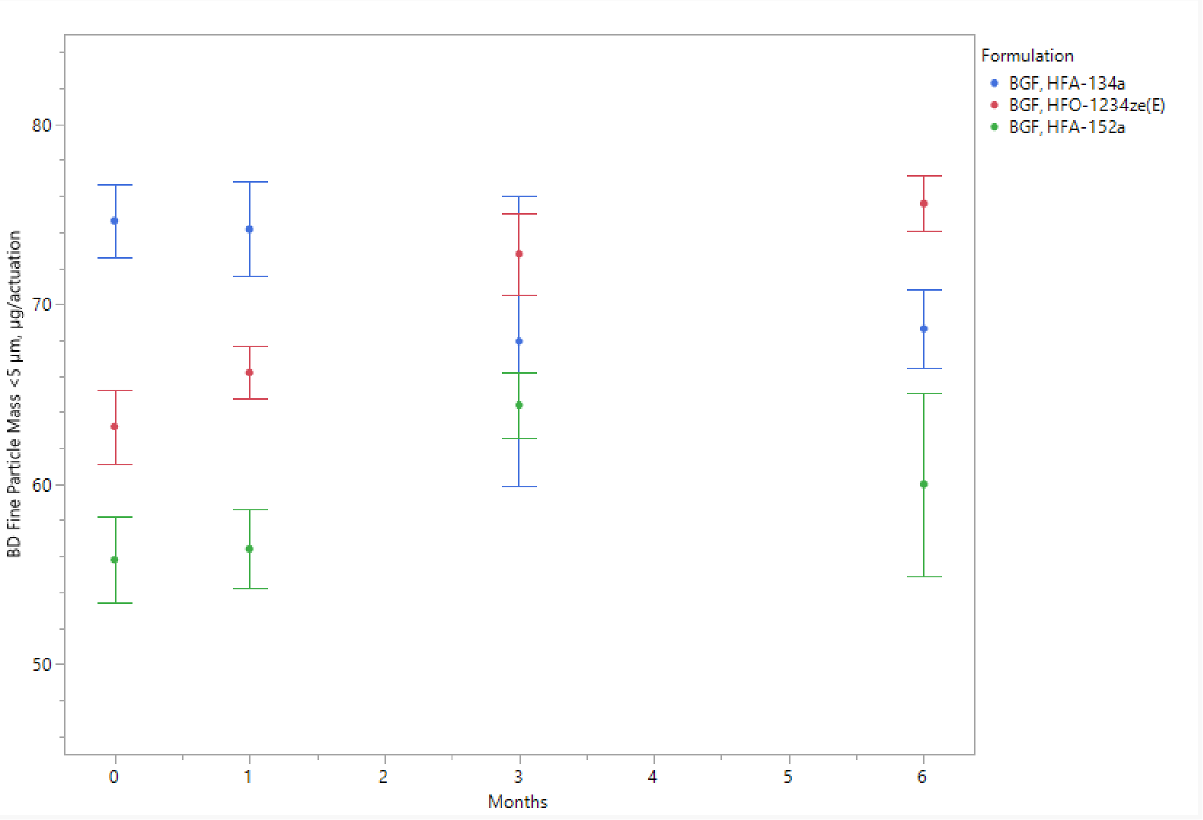


Results are presented as mean ± standard deviation.

BD, budesonide; BGF, budesonide, glycopyrrolate and formoterol fumarate; HFA, hydrofluoroalkane; HFO, hydrofluoroolefin; RH, relative humidity

Supplemental figure 13. Aerosol: fine particle mass < 5µm, expressed as micrograms/actuation, formoterol fumarate in each propellant under long-term stability conditions (25°C/60% RH) over 18 months

A

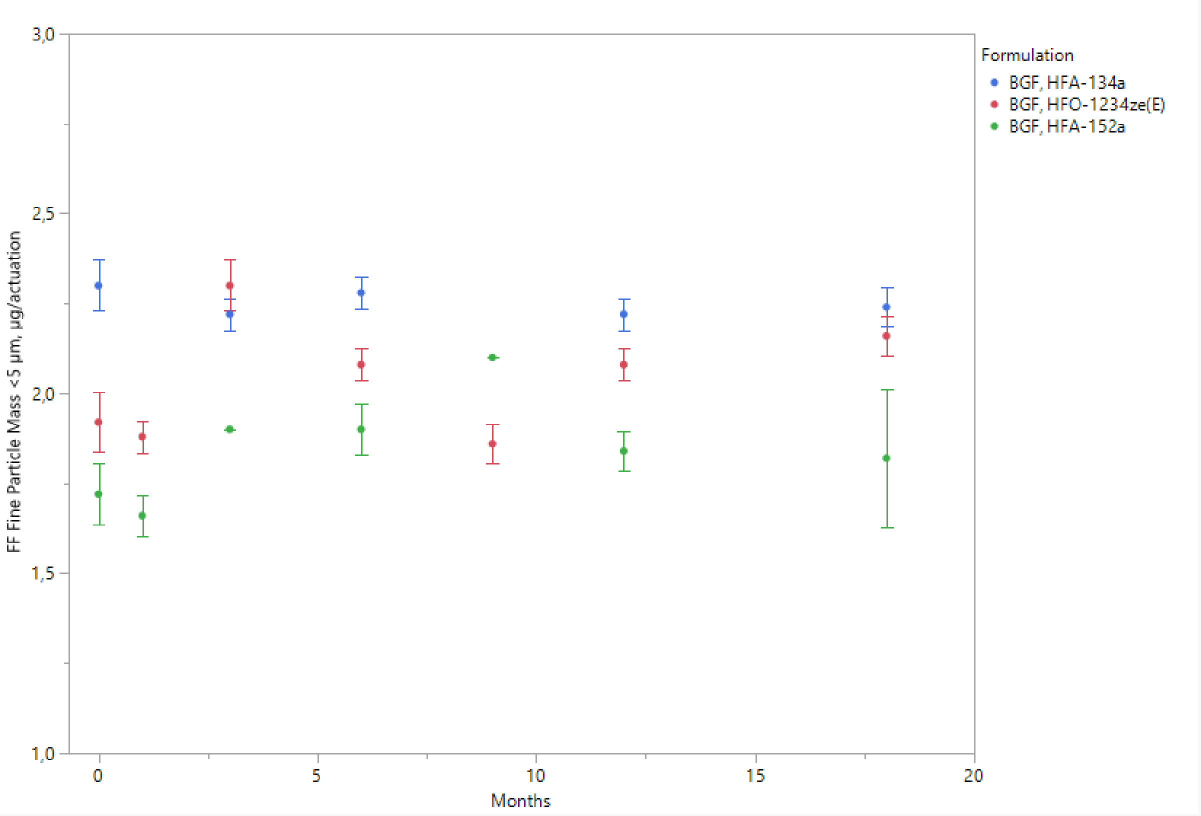


Results are presented as mean ± standard deviation.

BGF, budesonide, glycopyrrolate and formoterol fumarate; FF, formoterol fumarate; HFA, hydrofluoroalkane; HFO, hydrofluoroolefin; RH, relative humidity

Supplemental figure 14. Aerosol: fine particle mass < 5 µm, expressed as micrograms/actuation, for glycopyrrolate in each propellant under long-term stability conditions (25°C/60% RH) over 18 months

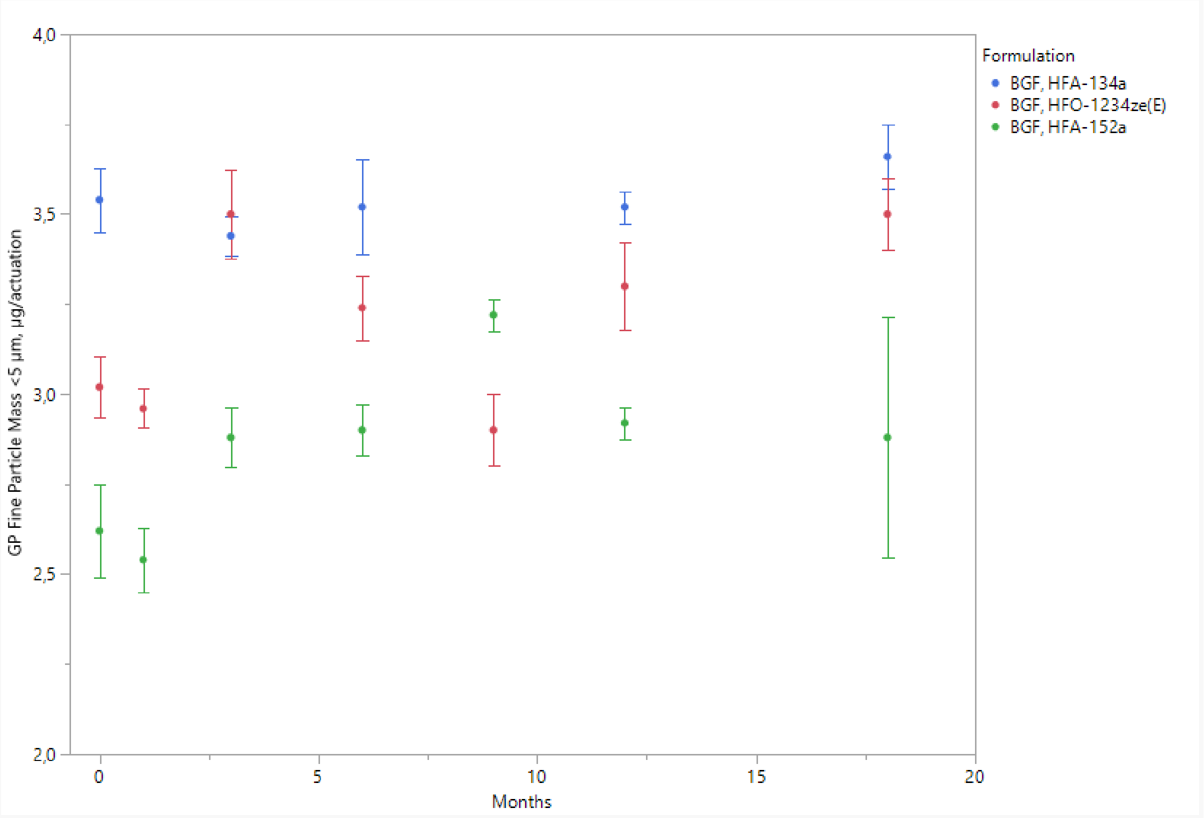


Results are presented as mean ± standard deviation.

BGF, budesonide, glycopyrrolate and formoterol fumarate; GP, glycopyrrolate; HFA, hydrofluoroalkane; HFO, hydrofluoroolefin; RH, relative humidity

Supplemental figure 15. Aerosol: fine particle mass < 5 µm, expressed as micrograms/actuation, for budesonide in each propellant under long-term stability conditions (25°C/60% RH) over 18 months

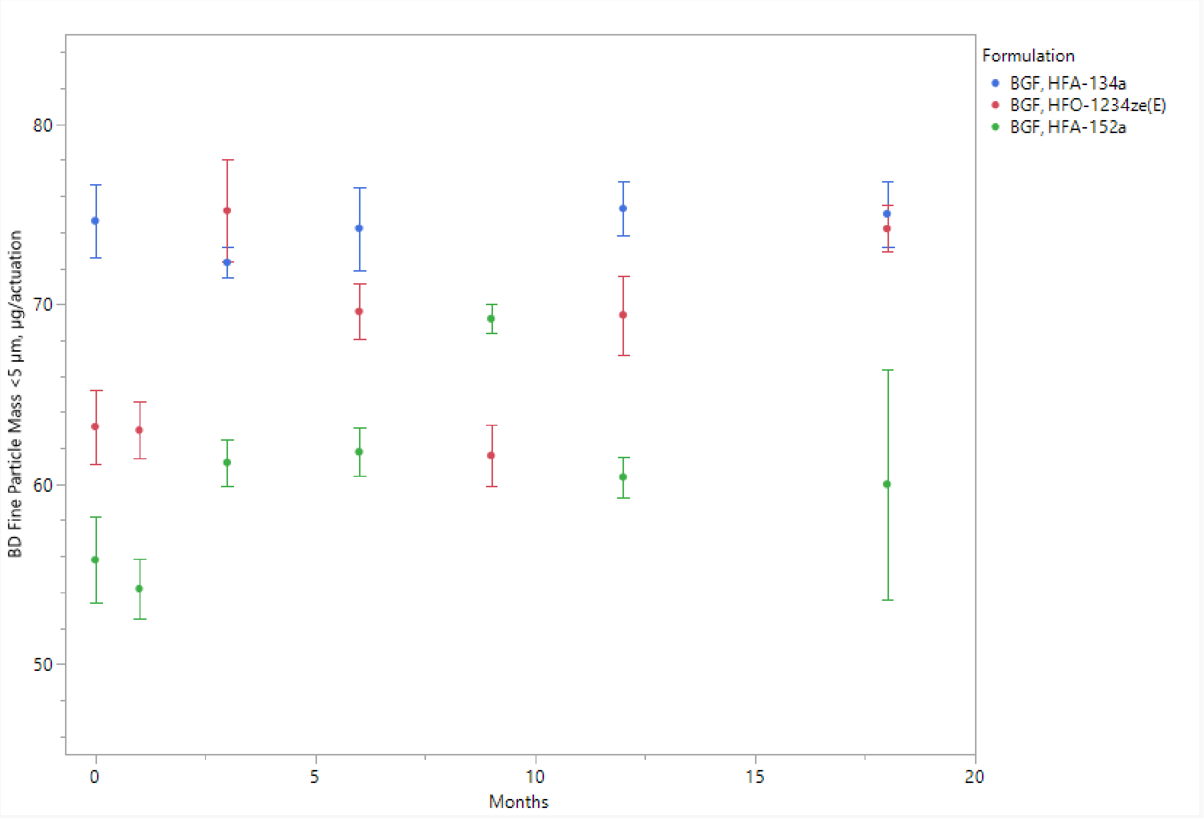


Results are presented as mean ± standard deviation.

BD, budesonide; BGF, budesonide, glycopyrrolate and formoterol fumarate; HFA, hydrofluoroalkane; HFO, hydrofluoroolefin; RH, relative humidity

Supplemental figure 16. Aerosol: fine particle fraction < 5µm, expressed as a percent of the label claim, for formoterol fumarate, glycopyrrolate and budesonide, in each propellant under accelerated stability conditions (40°C/75% RH) over 6 months

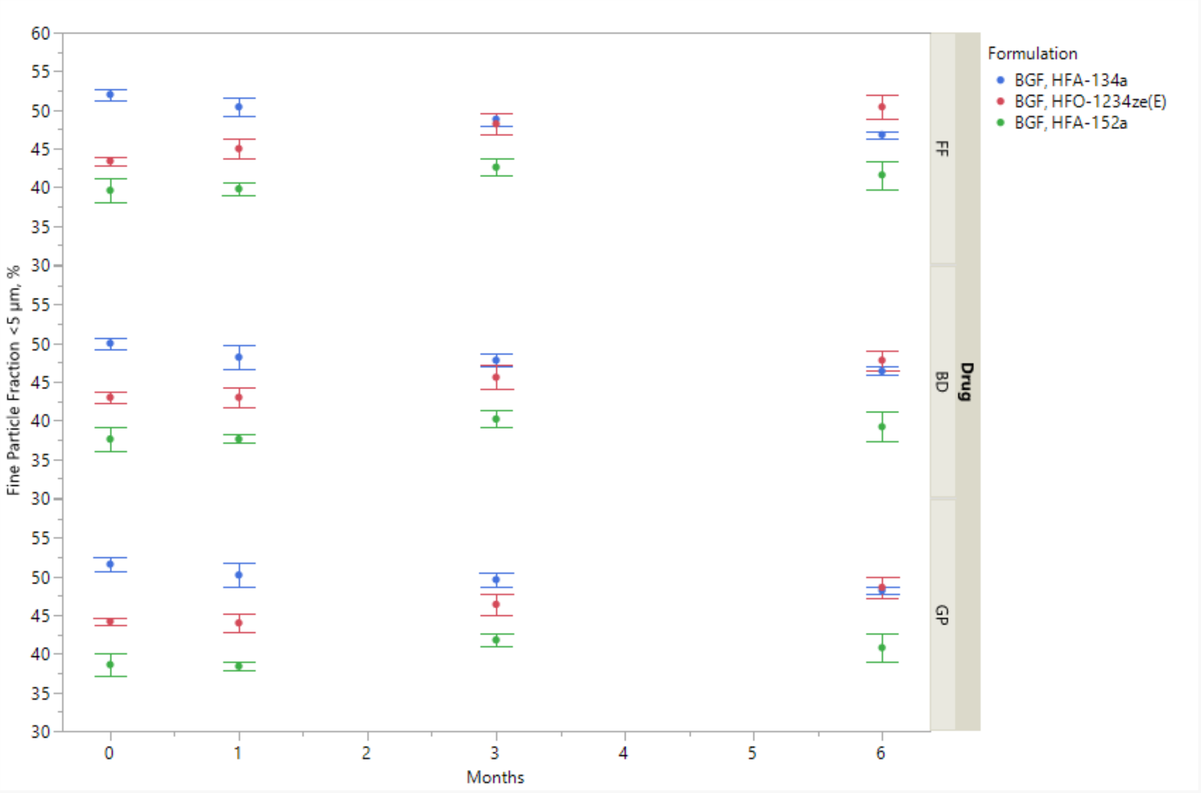


Results are presented as mean ± standard deviation.

BD, budesonide; BGF, budesonide, glycopyrrolate and formoterol fumarate; FF, formoterol fumarate; GP, glycopyrrolate; HFA, hydrofluoroalkane; HFO, hydrofluoroolefin; RH, relative humidity

Supplemental figure 17. Aerosol: fine particle fraction < 5µm, expressed as a percent of the label claim, for formoterol fumarate, glycopyrrolate and budesonide, in each propellant under long-term stability conditions (25°C/60% RH) over 18 months.

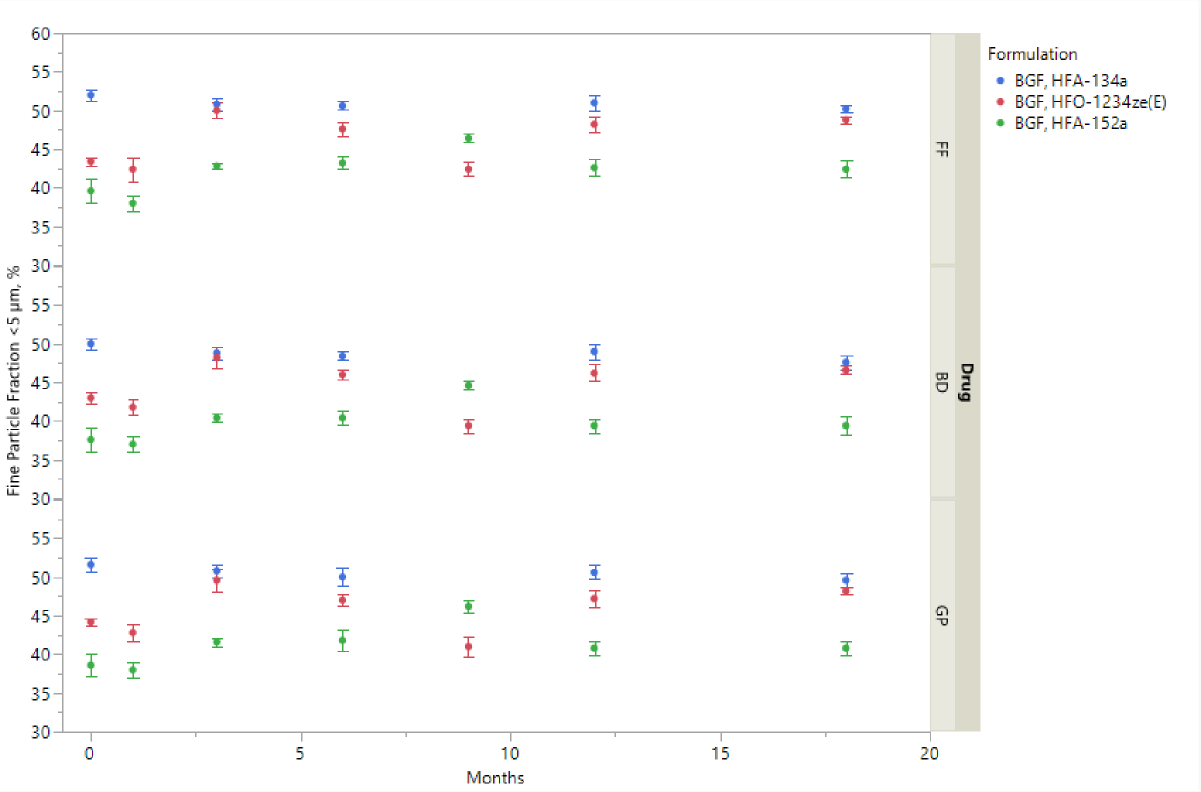


Results are presented as mean ± standard deviation.

BD, budesonide; BGF, budesonide, glycopyrrolate and formoterol fumarate; FF, formoterol fumarate; GP, glycopyrrolate; HFA, hydrofluoroalkane; HFO, hydrofluoroolefin; RH, relative humidity
